# Microbiome modulates immunotherapy response in cutaneous squamous cell carcinoma

**DOI:** 10.1101/2023.01.25.525369

**Authors:** Anita Y. Voigt, Andrew Walter, Timothy Young, John P. Graham, Bruna M. Batista Bittencourt, Alvaro de Mingo Pulido, Karol Prieto, Kenneth Y. Tsai, John P. Sundberg, Julia Oh

## Abstract

The gut microbiome is increasingly recognized to alter cancer risk, progression, and response to treatments such as immunotherapy, especially in cutaneous melanoma. However, whether the microbiome influences immune checkpoint inhibitor (ICI) immunotherapy response to non-melanoma skin cancer has not yet been defined. As squamous cell carcinomas (SCC) are in closest proximity to the skin microbiome, we hypothesized that the skin microbiome, which regulates cutaneous immunity, might affect SCC-associated anti-PD1 immunotherapy treatment response. We used ultraviolet radiation to induce SCC in SKH1 hairless mice. We then treated the mice with broad-band antibiotics to deplete the microbiome, followed by colonization by candidate skin and gut bacteria or persistent antibiotic treatment, all in parallel with ICI treatment. We longitudinally monitored skin and gut microbiome dynamics by 16S rRNA gene sequencing, and tumor burden by periodic tumor measurements and histologic assessment. Our study revealed that antibiotics-induced abrogation of the microbiome reduced tumor burden, suggesting a functional role of the microbiome in non-melanoma skin cancer therapy response.

## INTRODUCTION

With increasing recognition of the central role of the microbiome in infection, immune response, and inflammation, the microbiome’s role in cancer progression and treatment response is under extensive investigation ^1,2^. Occurrence of many cancers such as colon, gastric, head and neck cancers, and melanoma have been examined for microbial influences, and gut microbiome alterations have been linked to the efficacy of numerous cancer treatments, in particular the newer classes of immune checkpoint inhibitors (ICIs) targeting PD1 (programmed cell death protein 1, CD279) and CTLA4 (cytotoxic T-lymphocyte-associated protein 4, CD152) T cell receptors ^3-7^.

Melanoma has long been considered the most serious type of skin cancer with considerable metastatic potential and mortality. Nonetheless, non-melanoma skin cancers (NMSCs), such as squamous cell carcinoma (SCC) and basal cell carcinoma (BCC), are far more common, with high population incidence and a huge health care burden. Excessive exposure to ultraviolet radiation (UV) is the most common risk factor of NMSC, which is also tightly linked to other risk factors such as sex (men are more prone due to habits and occupations with increased UV exposure), baldness, and fair skin, the latter two providing reduced protection against UV radiation. Moreover, additional (microbiome-related) risk factors such as viral infections, immunosuppression (e.g., due to organ transplants), and chronic inflammation can increase susceptibility to skin cancer ^8-10^ and suggest the skin microbiome as a potential contributor to disease incidence, progression, and treatment outcome.

Most studies to date have focused on the gut microbiome and its interactions with cancer progression and treatment, but we hypothesized that the skin microbiome, which is a major influence on cutaneous immunity, may also impact skin cancer progression and treatment response. SCC develops from epidermal keratinocytes. Neoplastic cells are in close proximity to the epidermal surface and skin microbiome. Skin microbes indeed interact closely with specialized cells that express pattern-recognition receptors and can modulate immune and inflammatory response (i.e., T-lymphocyte function) ^11-13^.

We and others have found skin microbiome alterations in SCC, predominantly manifesting as a concomitant depletion of the potentially protective *Cutibacterium acnes* together with enrichment of *Staphylococcus aureus*, a skin pathobiont ^14-17^. Correspondingly, first insights have been identified into how microbes may affect SCC progression, for example by induction of inflammatory pathways by organisms such as *S. aureus* ^18^.

PD1 blockade can be highly effective against advanced melanoma, but this efficacy has been shown to be modified by the gut microbiome composition ^3-7,19,20^. Specific causative microbes differ between studies, but *Bacteroides* species, specifically *Akkermansia muciniphila*, are emerging as microbes of interest with therapeutic potential. Immunotherapy is used in patients with advanced SCC that cannot be treated with surgery or radiation therapy (e.g. Cemiplimab, a PD1 inhibitor) and PD1 blockade has been shown to delay SCC development in mice ^21^. But little is known about the potential modification of its efficacy by either the skin and/or gut microbiota. Further investigation is needed to begin to understand this potential new risk factor.

Here, we hypothesized that the skin and gut microbiome can modify the efficacy of immunotherapy in SCC. We selected representative microbes of the skin and gut microbiome that have been described to have antitumorigenic and immunostimulatory effects, such as *C. acne*s ^22-26^ and *A. muciniphila* ^6,19^. SCC tumors were induced in hairless mice via UV-B (ultraviolet B) radiation, and upon tumor development, ICI and bacteria or antibiotics – to test the effects of microbiome depletion – were administered. Tumor development and microbiome dynamics were monitored. Results showed that depletion of the microbiome with antibiotics, rather than the application of skin or gut bacterial candidates, improved the effect of immunotherapy.

## MATERIALS AND METHODS

### Animals

The albino hairless SKH1 mice (Crl:SKH1-*Hr*^*hr*^; Strain Code 477, abbreviated as “SKH1 mice”) were obtained from Charles River Laboratories, Wilmington, MA, USA. All mice were housed, and experiments performed in the animal facility at The Jackson Laboratory, Bar Harbor, ME, USA. All experiments were approved by The Jackson Laboratory Animal Care and Use Committees and performed in accordance with National Institutes of Health regulations. Please see Supplementary Materials and Methods for details.

### UV-B irradiation

SKH1 mice (51 mice: 46 females, 5 males) were 6-8 weeks of age at the initiation of UV-B light irradiation. The gender imbalance is a result of the sudden deaths of SKH1 mice (see Supplementary Materials and Methods). Mice were UV-B-irradiated Monday, Wednesday, and Friday for a total of 13 weeks with a total dosage of 750 mJ/cm^2^ for two weeks and a total of 1250 mJ/cm^2^ for the subsequent 11 weeks. Please see Supplementary Materials and Methods for details.

### Tumor measurements

Lesions were measured with a caliper, counted, and documented on a weekly basis and at the end point to assess the tumor burden. Animals with at least one lesion ≥4 mm were enrolled for antibiotics treatment, bacterial application, and immunotherapy treatment. Please see Supplementary Materials and Methods for further information.

### Antibiotics treatment

Once mice developed at least 1 lesion of ≥4 mm, mice were treated with topical and systemic antibiotics for 1 week (Group 1 and 2) prior to bacterial application and 4 weeks total in the case of Group 3. Please see Supplementary Materials and Methods for detailed information

### Microbial application

Group 1 received topical application of *C. acnes*, Group 2 received oral gavage of *A. muciniphila* while Group 3 received a long-term antibiotics treatment as well as a vehicle control topically applied and gavaged. Microbial application/gavaging recurred three times per week for 3 weeks after ICI/isotype administration until harvest. Details are described in the Supplementary Materials and Methods.

### Immunotherapy administration

In parallel to the bacterial treatment, mice received either the anti-PD1 monoclonal antibody or the corresponding isotype control. Mice were injected intraperitoneally three times per week for 3 weeks before bacterial administration. Details are described in the Supplementary Materials and Methods.

### Skin swab and fecal pellet collection and assessment of microbiome dynamics

Skin and gut microbiome dynamics were observed using skin swabs and fecal pellets, respectively. Swabs and pellets were collected prior antibiotics treatment, after antibiotics treatment and at the end point. DNA was extracted and subjected to 16S rRNA gene sequencing and analysis. Samples were also subjected to quantitative polymerase chain reaction to further illustrate depletion of the microbiome. All details are described in the Supplementary Materials and Methods.

### 16S rRNA gene data processing and analysis

An in-house civet pipeline using published tools was used to quality screen, data cleaning, and classify the sequencing data. All taxonomic features with a mean relative abundance of <0.001% and a prevalence of <10% were removed from the analysis. All details are described in the Supplementary Materials and Methods.

### Transplantable SKH1 tumor model

An immortalized cell line was generated by the Tsai Lab (Moffitt Cancer Center, Tampa, Florida, USA) from keratinocytes extracted from a histologically confirmed SCC tumor induced using UV-B light in SKH1 mice. Mice were intradermally injected with 250k cells (clone B610K), and tumors were measured on a weekly basis. Please see Supplementary Materials and Methods for detailed information.

### Tissue harvest and histology

At the endpoint, mice were euthanized and all skin lesions including adjacent normal appearing skin were dissected. The tissue was fixed in neutral buffered formalin, paraffin embedded, and H&E stained using standard protocols. Lesions were scored blindly by our histopathologist using published criteria ^27,28^. Please see Supplementary Materials and Methods for further information.

### Statistical analysis

R (version 4 ^29^) and RStudio (version 1.4.94) were used for statistical analysis and data visualization. Linear regression models were fitted, and repeated measure ANOVA was used to test statistical significance in tumor burden between two treatment groups over time. P-values were post-hoc multiple-test corrected using the Benjamini-Hochberg method and p-values <0.05 were deemed significant. Differences in tumor severity based on numbers of invasive and premalignant lesions confirmed by histologic evaluation were assessed using Fisher’s exact test. Please see Supplementary Materials and Methods for further information.

## RESULTS

### Study design

Per our previous study ^30^, we used ultraviolet B (UV-B) radiation to induce cutaneous squamous cell carcinoma (SCC) in albino hairless, inbred Crl:SKH1-*Hr*^*hr*^ (hereafter, SKH1) mice a commonly used mouse model used for dermatological and UV light-induced skin cancer studies ^27^. Fifty-one mice were UV-B irradiated for 13 weeks, three times 250 mJ/cm^2^ per week. Once tumors of ≥4 mm developed (defined at week 0), we randomized these mice into 3 groups to test the effect of candidate bacteria or microbiome depletion on tumor development. Each group was then subdivided into subgroups A and B (Table 1, Figure 1).

**Table 1:** Study design.

| Group | Bacteria | ICI | Abx | Sex<sup>^</sup> |
| --- | --- | --- | --- | --- |
| **Group 1A** | *C. acnes* | + | + | 8f |
| **Group 1B** | *C. acnes* | - | + | 9f |
| **Group 2A** | *A. muciniphila* | + | + | 7f, 2m |
| **Group 2B** | *A. muciniphila* | - | + | 8f, 1m |
| **Group 3A** | *Vehicle ctrl* | + | + * | 7f, 1m |
| **Group 3B** | *Vehicle ctrl* | - | + * | 7f, 1m |

\* 4 weeks of antibiotics (Abx) treatment (instead of 1 week), A: anti-PD1, B: Isotype control, ICI: Immunotherapy, Abx: Antibiotics, <sup>^</sup>Unbalanced due to unforeseen circumstances requiring adjustments to the experiment, see Material and Methods for details.

**Figure 1:**
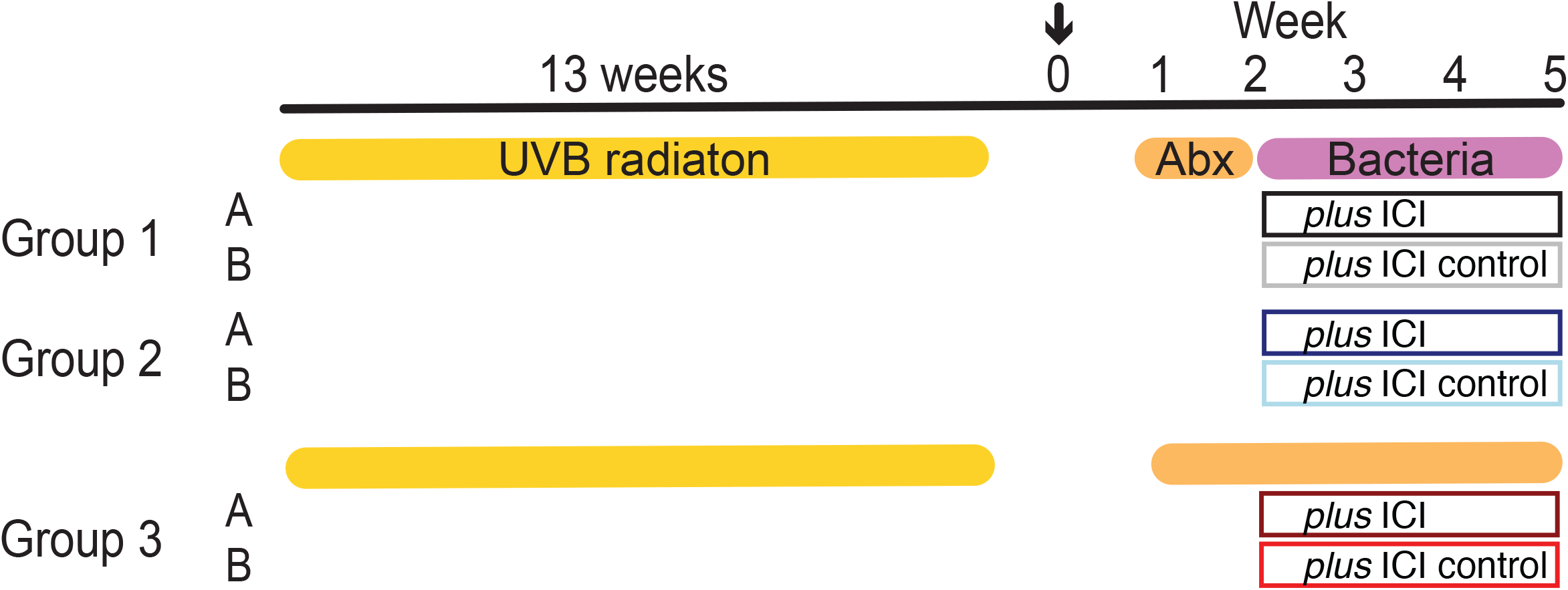
Experimental design. Mice were irradiated for 13 weeks, and upon tumor growth ≥4 mm, mice were assigned to a group and treated for 1 week (Group 1 and 2) with topical and systemic antibiotics followed by 3 weeks of bacterial application and ICI (Groups 1A and 2A) or isotype (Groups 1B and 2B) or 4 weeks with antibiotics and ICI or isotype (Group 3A and 3B). Week 0: enrollment (arrow), Week 1-5: antibiotics, bacteria and ICI treatment till end point. Skin and fecal samples were collected at weeks 1, 2, and 5. Antibiotics were applied topically and systemically. Details are described in the Supplementary Materials and Methods. Table 1 shows the Groups and animal numbers in more detail. ICI: Immunotherapy; UV-B: ultraviolet B; Abx: Antibiotics.

To deplete the resident flora, all groups received one week of broadband antibiotics treatment. Subsequently, Group 1 received topical application of *C. acnes*, Group 2 received orally gavaged *A. muciniphila*, while Group 3 continued antibiotic treatment for the duration of the experiment (4 weeks total) and received topically applied and orally gavaged vehicle control. Bacteria and vehicle control were applied twice per week for 3 weeks. Of each group, subgroup A received anti-PD1 immunotherapy (anti-PD1), while subgroup B received the corresponding isotype control for 3 weeks (weeks 2-5, Figure 1).

To monitor skin microbiome composition and load, skin swabs and fecal samples were collected at the following timepoints: just before antibiotics treatment (week 1), after one week of antibiotics treatment (week 2), and at end point three weeks later (week 5). At the end point, two skin swab samples were collected per mouse: a collective swab of the healthy, tumor-adjacent skin and a collective swab of all lesions to monitor SCC-associated microbiome dynamics. Samples were subjected to 16S rRNA gene sequencing to reconstruct microbiome composition, and qPCR to determine antibiotic depletion. More details can be found in the Material and Methods.

### Microbial load and dynamics upon antibiotic treatment and microbial engraftment

Cutaneous and fecal microbiome were depleted with a 1-week (Group 1 and 2) or 4-weeks (Group 3) topical (Mupirocin and Neosporin) and systemic (ampicillin, streptomycin, vancomycin and colistin) antibiotics course. We used read counts from 16S rRNA gene sequencing as a rough proxy for biomass to illustrate the change of the microbiome induced by the antibiotics treatment and administration of *C. acnes* and *A. muciniphila* (Figure 2, S1 and S2). As expected, the microbiome biomass declined 1 week after treatment in all groups and was most pronounced in stool rather than on the skin (Figure S1). Biomass increased over time in Groups 1 and 2 after cessation of antibiotics treatment (due to natural re-colonization), and the skin and feces showed effective engraftment of the topically applied or gavaged *C. acnes* and *A. muciniphila* (Figure S2). In group 3, microbial biomass in skin showed further declines until the endpoint, likely due to the persistent topical and systemic antibiotics treatments. In the gut, the decline was consistent between week 2 and 5 in Group 3A, but with a modest increase in *Clostridiales* (potentially resistant to the antibiotics) at the end point in group 3B (Figure S2). We further performed 16S qPCR of DNA samples before (week 1) and after (week 2) antibiotics to verify depletion, as read counts are only a proxy for microbial abundance and can be skewed by sequencing artifacts. Here, we measured the fold change of the 16S rRNA from before to after, which also emphasized a large decline in 16S rRNA gene DNA after antibiotic treatment (Figure 2). Overall, we concluded that the topical and systemic antibiotics sufficiently reduced microbiome biomass both in short term and facilitated engraftment of the administered microbes (Group 1 and 2) or suppressed the microbiome persistently.

**Figure 2:**
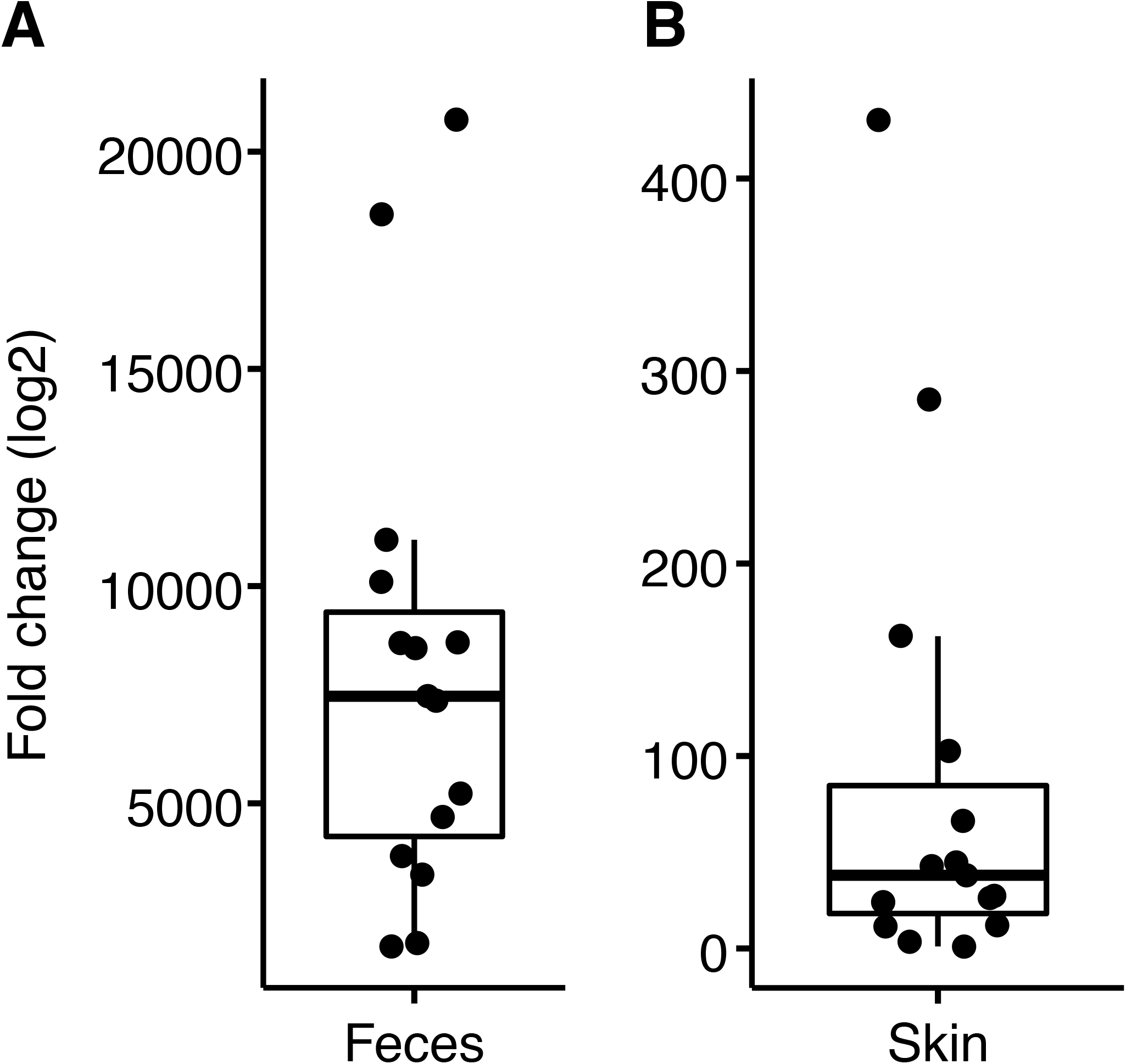
Bacterial depletion by concurrent topical and systemic antibiotic treatment. qPCR targeting the 16S rRNA gene was performed on each sample (in triplicate) to compare before versus after antibiotics treatment (Supplementary Materials and Methods). A representative set of paired **A**: fecal (n=15) and **B**: skin (n=15) samples were assessed for their microbial content before and after antibiotics treatment using qPCR. Fold change (log2) of before versus after antibiotics treatment is shown to contextualize reduction in microbial biomass.

### UV-induced tumor severity and burden depends on treatment

Tumor burden of each mouse was assessed in two ways: 1) histologic evaluation of each harvested lesion (>2mm) and 2) weekly caliper measurements and subsequent calculation of a panel of parameters according to ^28^: tumor volume, 2D tumor area, multiplicity (average number of tumors per mouse) and Knud Thomsen (KT) Three-Dimensional (3D) Surface (apportions optimal weightage to the three tumor dimensions) size. Table 2 shows a summary of all histologic observations (full data, Table S1) where fully and microinvasive tumors were combined as “invasive SCC” and preneoplastic and premalignant tumors were aggregated as “premalignant SCC”. Exophytic papilloma and fibropapilloma were summarized as “papilloma”. Based on the histologic evaluations, the total number of diagnosed invasive SCC lesions detected across all mice was higher in the Group 1 and 2 mice than Group 3, irrespective of ICI treatment (Table 2). However, sub-categorizing lesions (i.e., SCC vs. no SCC, premalignant vs. no premalignant, papilloma vs. no papilloma), showed no significant difference (Fisher’s exact test, p>0.05 for all comparisons). Particularly for SCC, this indicated that the Groups 1 and 2 mice with SCC were likely to have multiple SCCs, rather than a higher prevalence of SCC. We also performed immunohistochemistry for select immune markers to characterize the immunological response to the different treatments. We concluded that our observations were rather an effect of the hairless mutant skin phenotype than the treatment (more details in Material and Methods and Supplementary Material and Methods).

**Table 2:** SCC-related histologic observations. Full list of observations is in Table S1.

|  | Group |  |  |  |  |  |
| --- | --- | --- | --- | --- | --- | --- |
|  | 1A | 1B | 2A | 2B | 3A | 3B |
|  | 8 mice | 9 mice | 9 mice | 9 mice | 8 mice | 8 mice |
| **Invasive SCC\*** | 10 | 11 | 13 (12) | 13 (9) | 5 (5) | 1 (1) |
| **Premalignant SCC#** | 40 | 30 | 32 (23) | 22 (20) | 23 (19) | 28 (22) |
| **Papilloma“** | 0 | 1 | 1 (1) | 0 (0) | 0 (0) | 0 (0) |

\* fully and microinvasive SCC tumors; # all preneoplastic/pre-malignant observations; “ Exophytic papilloma/ fibropapilloma; Numbers in brackets are observations for female mice only for the groups that included 1-2 male mice.

While the histologic analysis showed no significant difference in tumor severity between groups, we observed differences in tumor growth kinetics over time as measured by KT-3D Surface area and tumor volume (Figure 3) and 2D tumor area and tumor multiplicity for all mice (irrespective of sex, Figure S4). Due to the unbalanced design of predominantly female and few male mice (Table 1), we calculated statistical tests for all mice and for females only (Table S4), highlighting results for “females only” when discordant.

**Figure 3:**
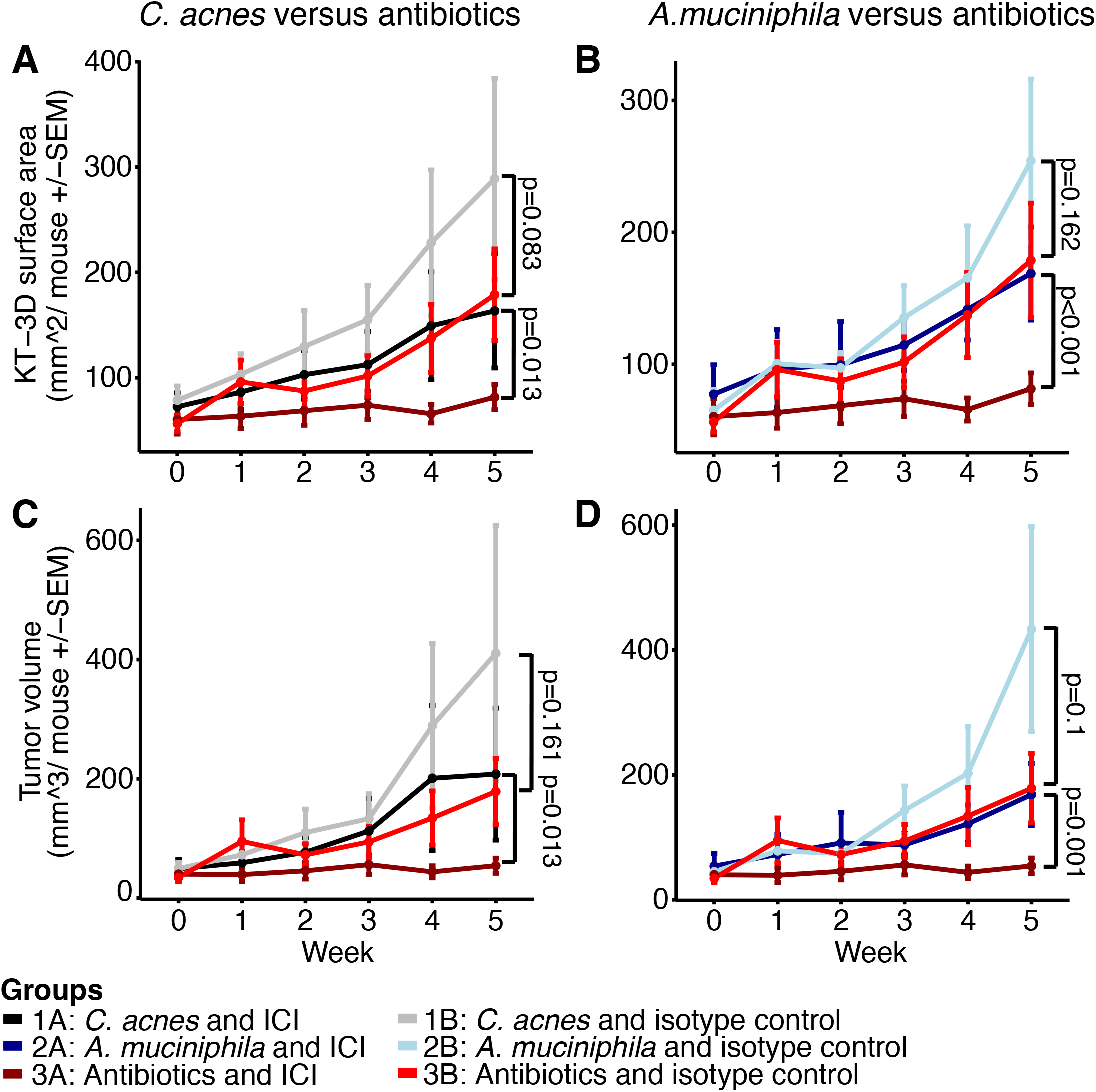
Tumor surface area and volume. **A** and **C**: Depicted are the KT3D tumor surface area and **B** and **D** tumor volume for the *C. acnes* (Group 1, **A, B**) and *A. muciniphila* (Group 2, **C, D**) treated groups compared to the antibiotic-treated mice (Group 3). Week 0 represents the first week of tumor measurement (enrollment), week 1 pre-antibiotics treatment and weeks 2-5 bacteria/antibiotics as well as ICI/isotype treatment with euthanasia at the end point (week 5). P-values were calculated using linear modeling and repeated measure ANOVA including the weeks 3, 4 and 5 (details in Supplementary Materials and Methods, full list of p-values in Table S4). Depicted are all mice per group irrespective of sex: Group 1A: n=8, Group 1B: n=9, Group 2A: n=9, Group 2B: n=9, Group 3A: n=8, Group 3B: n=8.

Examining the effect of ICI versus isotype treatment using KT-surface area, only Group 3 mice treated with ICI exhibited a significant reduction. In terms of tumor volume, a significant reduction was observed for Group 2 (not significant for females only) and Group 3, but not for Group 1. As expected, we observed a significant anti-tumorigenic effect of ICI on the 2D tumor area (for all groups) in mice subjected to ICI compared to mice treated with the isotype control. Tumor multiplicity was not affected by the ICI treatment, indicating that tumors, once measurable, did not resolve upon treatment. P-values for each comparison are listed in Table S4.

Surprisingly, comparing mice treated with ICI and bacteria (Groups 1A and 2A) to mice on the long-term antibiotics (Group 3A), revealed a significantly reduced tumor burden in terms of KT-surface, area, and volume, but not multiplicity, in Group 3. On the other hand, the comparison of mice treated with the isotype control and bacteria (Groups 1B and 2B) to Group 3B, did not result in a statistically significant difference apart from borderline significance for multiplicity for 1B versus 3B (possibly due to the large variability in 3B). Although the median of the tumor growth parameters for antibiotics-treated mice were generally lower than in groups treated with bacteria, the large variance observed rendered the statistical significance of these results borderline or non-significant. We note that such variance is inherent with the use of the outbred SKH1 mouse model and the UV-induced (spontaneous) skin cancer model. The lowest variability was observed in Group 3A, suggesting a synergistic effect of antibiotics and immunotherapy treatment. Taken together, these results indicated that antibiotic treatment had no effect on tumor growth kinetics when administered with the isotype control. However, when administered in combination with ICI, depletion of the microbiome enhanced its anti-tumorigenic effect.

### Transplantable SCC model confirms antitumorigenic effect of antibiotics

To mitigate the mouse-to mouse variability and labor-intensive induction of the outbred SKH1 SCC model, and to further test the effect of antibiotics on tumor development, we obtained an immortalized cell derived from an UV-induced tumor of a female SKH1 mice. We intradermally injected different numbers of cells (250k and 500k) to mimic the potentially different tumor burdens into antibiotic treated and untreated SKH1 mice (transplantable model, Material and Methods). We again performed weekly caliper measurements of the injected tumors and assessed treatment-related differences. Only the kinetics of tumor volume over time is shown due to the relatively homogeneous manifestation and growth of identically injected cells (Figure S5).

Male SKH1 mice generally exhibited accelerated tumor growth compared to females. This could potentially be explained by the derivation of the tumor cells from a female SKH1 mouse. No antibiotic-related differences were identified for mice injected with 500k cells or females injected with 250k cells. However, males injected with 250k cells and antibiotics treatment showed significantly reduced tumor volume compared to untreated mice (Wilcoxon rank-sum test: for week 3.5: p = 0.0005, for week 3: p=0.0147, repeated measures ANOVA: p<0.001, Figure S5). This supports our previous findings in which the long-term antibiotic treated mice exhibited a reduced tumor burden compared to mice on short-term antibiotics and supplemented with bacteria (Group 1 and 2) even if not always significant due to the large variability. It is noteworthy (but not unexpected) that larger amounts of transplanted cells (e.g., 500k cells) overpowered the effect size of antibiotic treatment.

## DISCUSSION

Here, we have shown that the growth of spontaneous SCC and a transplanted SCC cell line was reduced in a hairless, outbred mouse model upon manipulation of the microbiome. Long-term broadband antibiotics resulting in a reduction of the endogenous skin and gut microbiome enhanced the anti-tumorigenic effect of anti-PD1 ICI over short-term antibiotics treatment followed by colonization with a candidate skin (*C. acnes*) or gut (*A. muciniphila*) microbe. This was despite our selection of *C. acnes* based on its ascribed anti-tumorigenic activity ^22-26^ and *A. muciniphila* based on its a favorable impact on ICI response in other cancer types (e.g., ^6,19^).

In addition, previous reports (e.g., ^6^) showed that antibiotics typically compromised the effectiveness of the PD1 blockade in mouse tumor models and patients with advanced cancers. By contrast, we noted an improvement of the ICI efficacy with concomitant antibiotic use over the short-term antibiotic usage plus skin/gut microbial colonization. The discordance likely stems at least in part from 1) the use of different melanoma and sarcoma cell lines in a subcutaneous model of non-SKH1 mouse strains (C57BL/6 and BALB/c), 2) the study of cancer patients with advanced non-small cell lung cancer, renal cell carcinoma, or urothelial carcinoma, rather than our focus on SCC, and/or 3) our use of antibiotics that depleted both the gut and the skin microbiomes. These results indicate a specificity and the complexity inherent to microbial contribution to cancer progression and treatment response. We note that while we demonstrated effective compartmentalization of gut vs. skin microbial colonization, it is impossible to disentangle the effects of topical vs. systemic antibiotic treatment on skin cancer development (or on the skin or gut immune system) since both were administered in parallel.

In addition, we note inherent limitations of the outbred, spontaneous UV-induced SCC model. This is a labor-intensive model, given the lengthy irradiation needed to spontaneously induce SCC, which limits cohort size due to the high effort spent per mouse. Moreover, the inter-mouse variation in tumor growth kinetics and counts was large, which complicated initiation of the antibiotics and ICI treatment timing. Secondly, the histologic evaluations confirmed fully invasive SCCs in many mice including ICI-treated mice, indicating that the mice potentially had too advanced tumors at ICI onset for the ICI to be highly effective. Thus, we only observed decelerated tumor growth but no tumor shrinkage or reduction of tumor multiplicity upon treatment. The tumor burden in the spontaneous model likely superseded the effect size of the ICI treatment and any potential effect of the microbes applied. On the other hand, the administration of probiotics of defined microbial composition was detrimental to ICI treatment response via immune mechanisms in melanoma patients and a preclinical mouse model ^31^. Fine tuning of this model to identify a more appropriate tumor burden to evaluate microbial and ICI effects, as well as further deployment of the transplantable SCC model in which we showed that long-term antibiotics usage can delay the tumor growth (even without ICI) at moderate tumor burden as well as validation experiments in germ-free mice, would be beneficial in subsequent studies.

In summary, our study provides compelling justification for further investigations of the role of the skin and gut microbiome in SCC cancer progression and treatment response, to better characterize the potential of these bacteria as therapeutic targets and biomarkers of treatment response for this common cancer.

## ABREVIATIONS

NMSC: Non-melanoma skin cancer
SCC: Squamous Cell Carcinoma
OTU: Operational taxonomic unit
ICI: Immune checkpoint inhibitor
UV-B: ultraviolet B
PD1: programmed cell death protein 1
KT-surface: Knud Thomsen three-dimensional (3D) surface

## DATA AVAILABILITY

The datasets related to this article can be found in the Supplementary Material. Abundance tables used for taxonomic analysis are provided in Table S3.

## ACKNOWLEDGEMENTS AND FUNDING

We are thankful to the Oh group for inspiring discussions, Genome Technologies at The Jackson Laboratory for Genomic Medicine for support with sample processing and sequencing, Dr. Hoan Nguyen for support with sequence data processing, and the Histopathology facility at The Jackson Laboratory. We acknowledge the breeding performed by Jane Branca and the mouse experiments performed by the Gnotobiotic and Microbiome Mouse Program (GMMP) team (Keira Eschete, Amber Robbins, Breah Farrington, Ed Keniston, Dina Baker, Karen French) and veterinarians (Terri Iwata, Bonnie Lyons) as well as Beverly Macy in Bar Harbor and Omar Chavez Chiang at Moffitt Cancer Center. We thank Joseph Brown from the Microbial Genomic Services at The Jackson Laboratory for Genomic Medicine for support with sample processing. We are grateful for the funding of AYV through the Pyewacket PostDoc Fund. This research received further funding through the American Cancer Society (IRG-82-003-33 NCCC-01 and RSG-16-255-01), Scott R. MacKenzie Foundation, and the LEO Foundation. The Jackson Laboratory Shared Services were supported in part by Basic Cancer Center Core Grant from the National Cancer Institute (CA034196). JO is additionally supported by the NIH (1 R01 AR078634, DP2 GM126893 and K22 AI119231, 1U54NS105539, 1 U19 AI142733, 1 R21 AR075174).

## AUTHOR CONTRIBUTIONS

Conceptualization: AYV, JO, KYT, JPG, JPS, TY; Data curation: AYV, JO, JPS; Formal Analysis: AYV, JPS; Funding acquisition: JO; Investigation: AW, BMB, AMP, KP; Project Administration: AYV, JO, TY; Resources: JO; Supervision: JO, AYV, TY; Visualization and Writing: AYV, JO; Writing - Review and Editing: JO, AYV, KYT, JPS, TY, AW, JPG, BMB, AMP, KP.

---

**Figure S1:**
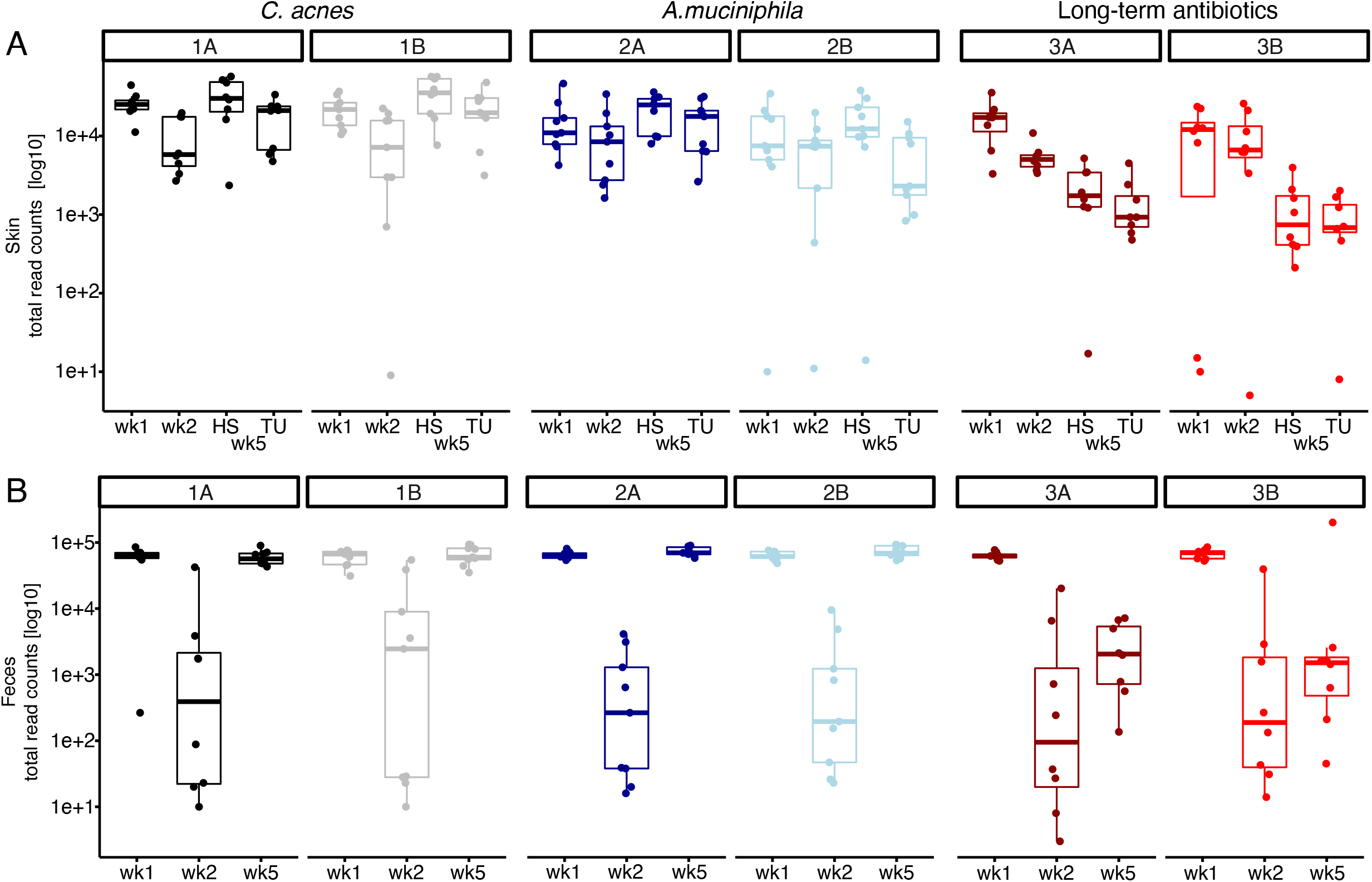
Raw read counts of 16S sequenced samples across time points and groups. Boxplots show total microbial read counts identified per sample in A: skin and B: fecal samples before (week 1, wk1) and one week after (week 2, wk2) antibiotics treatment as well as at the end point (week 5, wk5) for each group. HS: healthy skin and TU: tumor at endpoint. Metadata is listed in Table S2, and the Operational taxonomic unit (OTU) table is in Table S3. Depicted is the data for all mice irrespective of sex: Group 1A: n=8, Group 1B: n=9, Group 2A: n=9, Group 2B: n=9, Group 3A: n=8, Group 3B: n=8.

**Figure S2:**
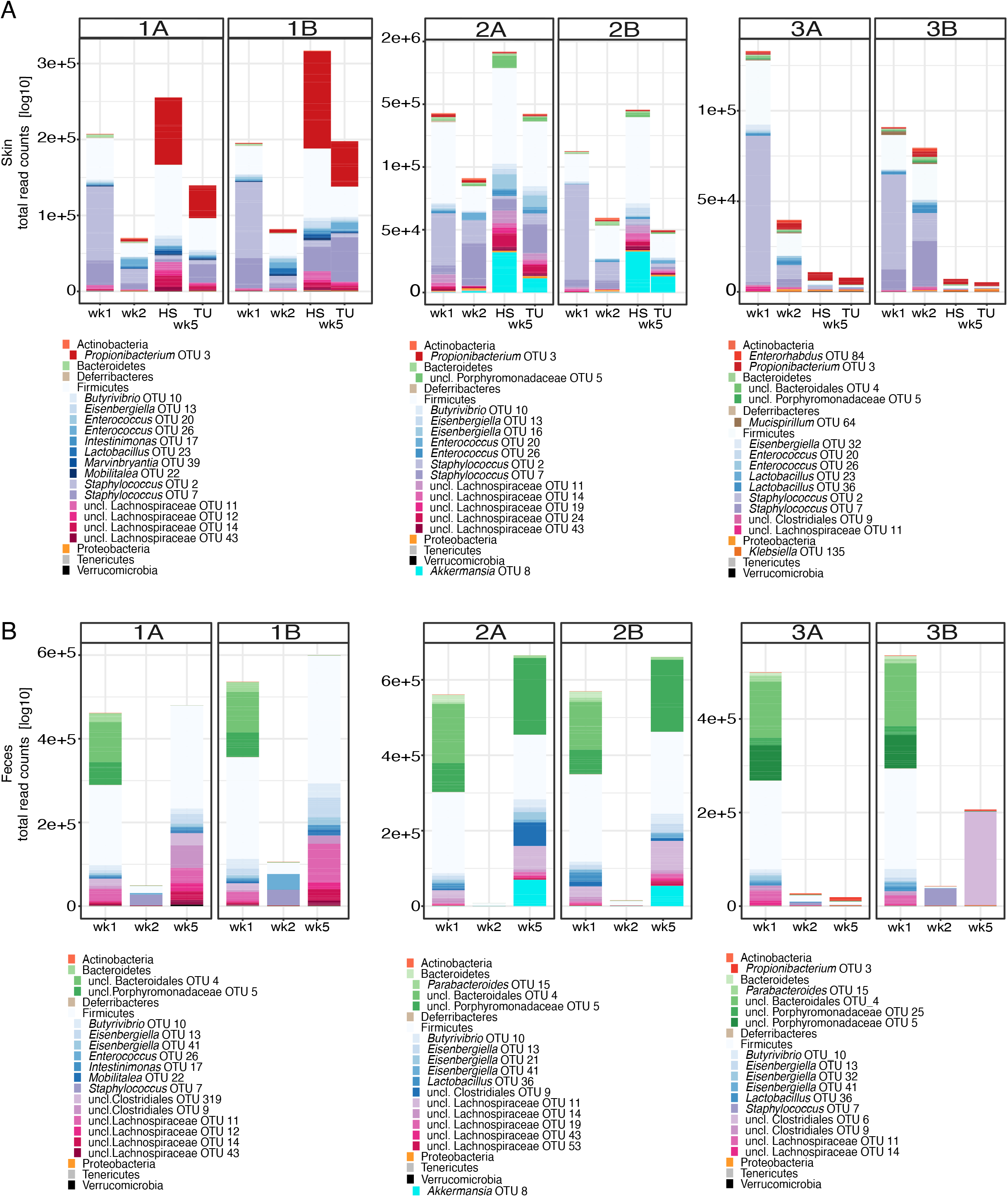
Estimated load of most abundant species, based on read counts as proxy for microbial biomass. Abundance plot A shows the microbial communities in skin and B in the fecal samples before (week1, wk1) and after one week (week2, wk2) of antibiotics treatment as well as at the end point (week 5, wk5). Microbes that were applied topically or gavaged in the respective groups were highlighted in red (*C. acnes*) and cyan (*A. muciniphila*). HS: healthy skin and TU: tumor at endpoint. Metadata is listed in Table S2 and the OTU table is in Table S3. Depicted is the data for all mice irrespective of sex: Group 1A: n=8, Group 1B: n=9, Group 2A: n=9, Group 2B: n=9, Group 3A: n=8, Group 3B: n=8.

**Figure S3:**
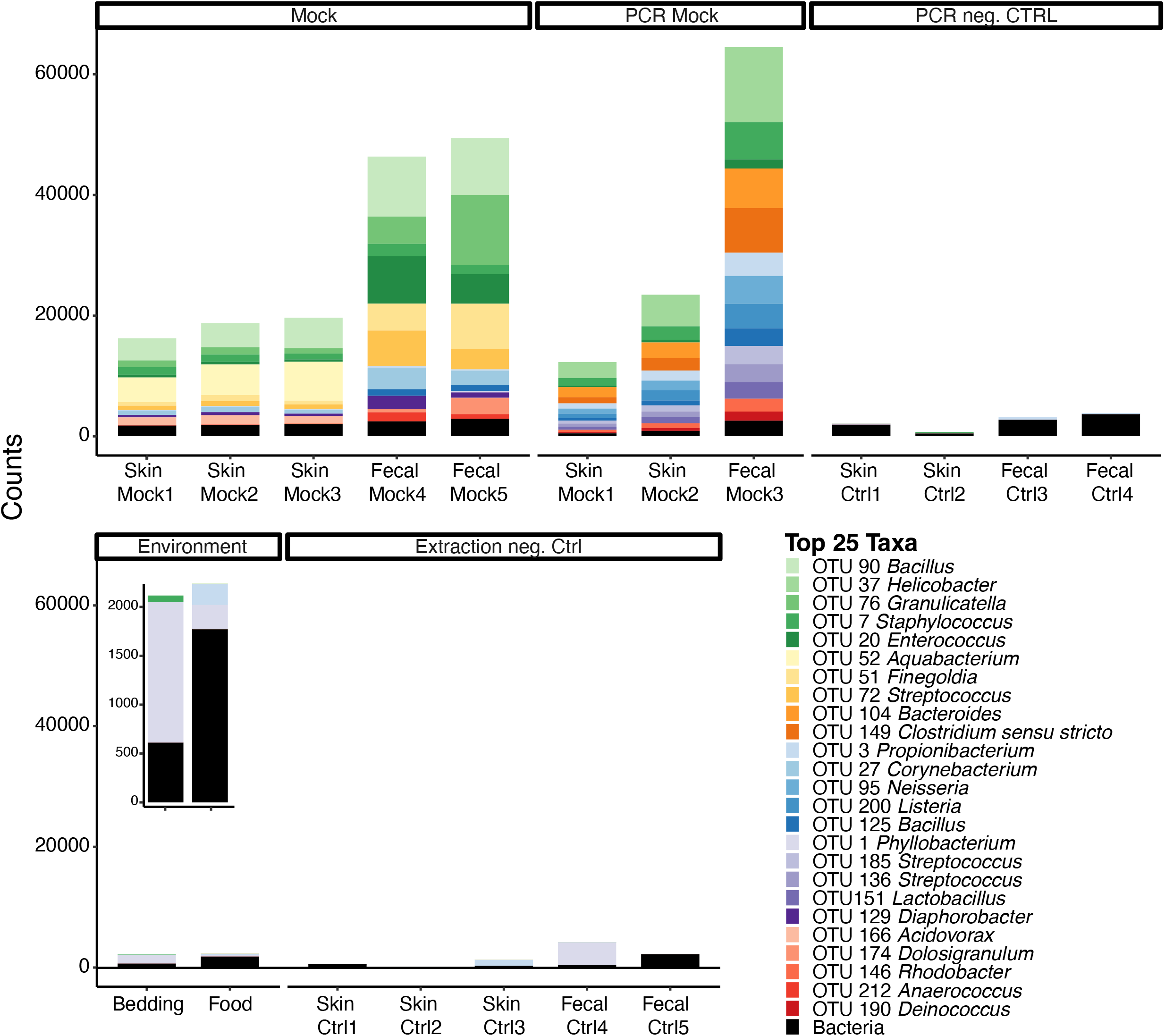
16S read counts plots for negative and positive controls. Depicted are the 25 most abundant OTUs identified in the control samples while the remainder OTUs are summarized as “Bacteria”. An in-house mock community (25 diverse Gram-positive bacteria, Gram-negative bacteria and fungi) was used as positive controls. The insert shows the enlarged counts plot for bedding and food. Metadata is listed in Table S2 and the OTU table is Table S3. OTU1, a *Phyllobacterium* was identified in bedding and food controls and removed from the dataset (see Material and Methods). *C. acnes* versus antibiotics

**Figure S4:**
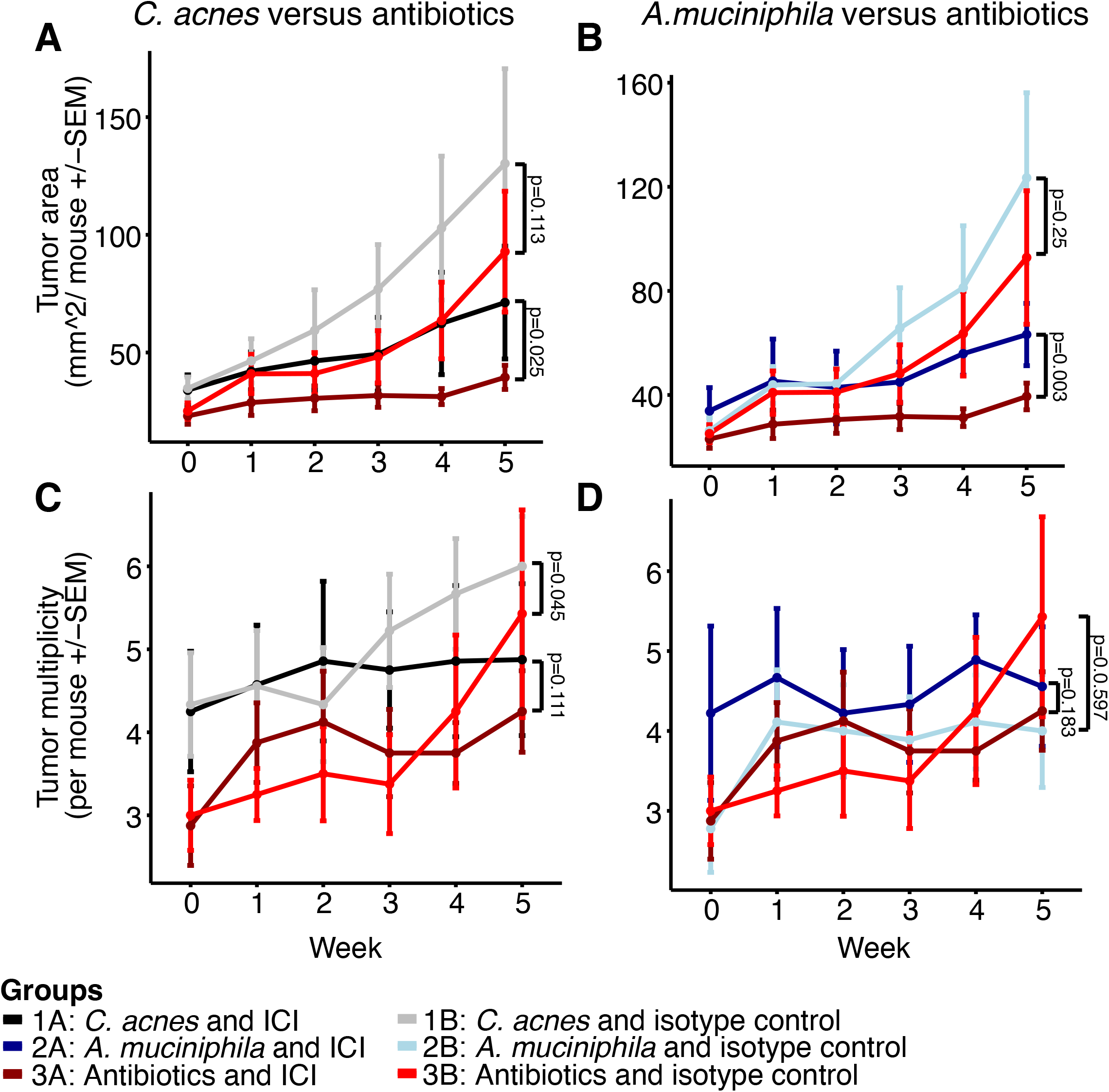
Tumor area and multiplicity. A and B: Depicted are the tumor area and (C and D) tumor multiplicity for the *C. acnes* (Group 1, A, C) and *A. muciniphila* (Group 2, B, D) treated groups compared to the antibiotics treated mice (Group 3). Week 0 represents the initial week of tumor measurement (enrollment), week 1 pre-antibiotics treatment and weeks 2-5 bacteria/antibiotics as well as ICI/isotype treatment with euthanasia in the final week (end point, week 5). P-values were calculated using linear modeling and repeated measure ANOVA including the weeks of treatment 3, 4 and 5 (details in Material and Methods and full list of p-values in Table S4). Data of all mice per group are shown irrespective of sex: Group 1A: n=8, Group 1B: n=9, Group 2A: n=9, Group 2B: n=9, Group 3A: n=8, Group 3B: n=8 (Table 1).

**Figure S5:**
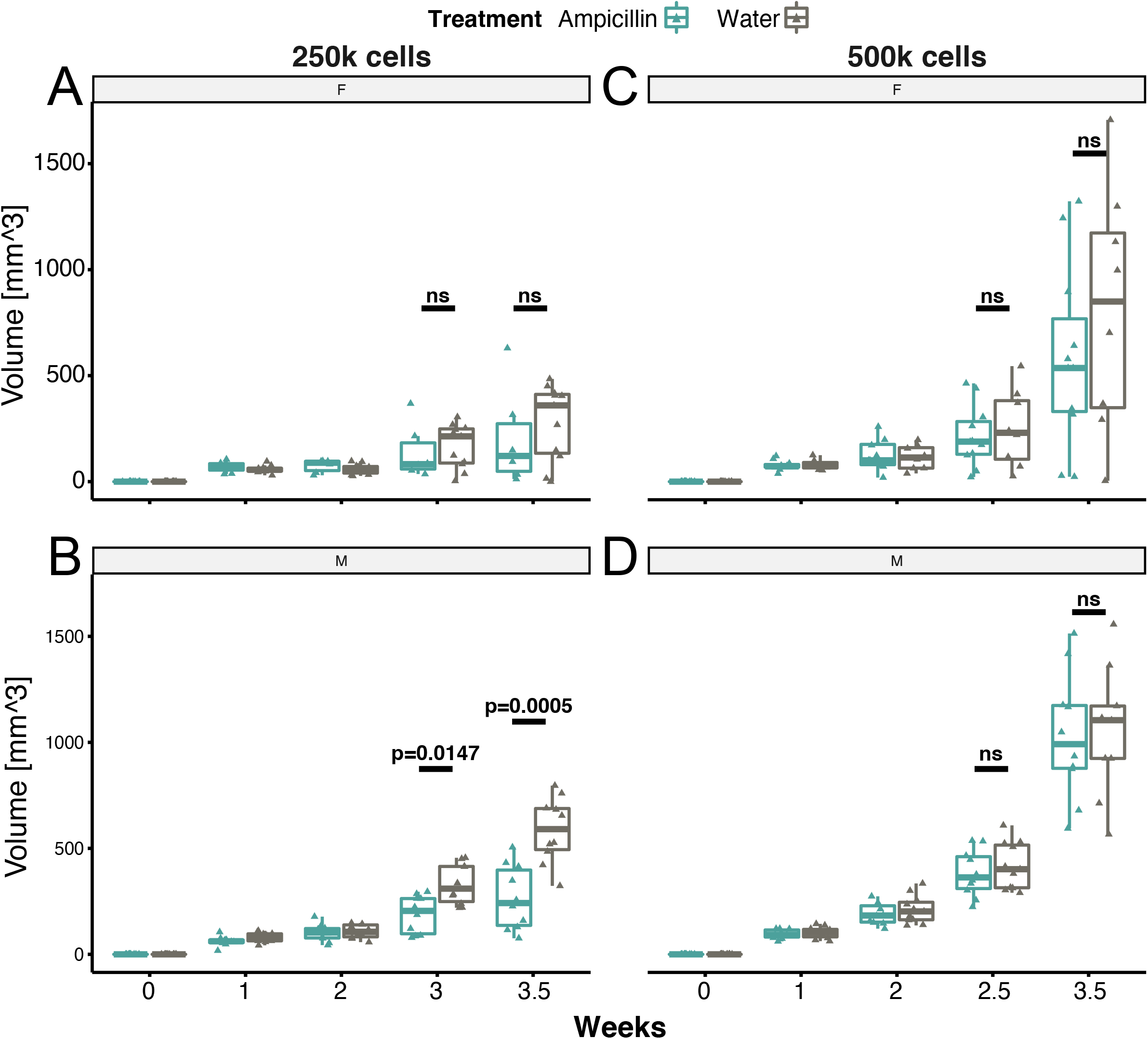
Tumor growth in transplantable model over time. Boxplots show the volume of the tumors transplanted into female (A and C) and male (B and D) mice injected with different cell numbers (250k and 500k per injection) with and without antibiotics. Tumor growth was significantly reduced in male mice treated with antibiotics versus untreated mice when 250k cells were transplanted. Group differences were assessed using repeated measure ANOVA. Initially, 8-11 mice per group and sex were used but a few mice dropped out during the course of the experiment due to adverse side effects (such as dehydration, detailed in Material and Methods). ns: not significant.

