## Supplemental Material and Methods for "Microbiome modulates immunotherapy response in cutaneous squamous cell carcinoma"

### CONTENT

1.) **Supporting Data**

2.) **Supplementary Material and Methods**

3.) **Supplemental Table Legends**

**Table S1:** Detailed list of histologic observations.

**Table S2:** Metadata.

**Table S3:** 16S read counts table.

**Table S4:** Table of all p-values calculated for each comparison.

4.) **Supplemental Figure Legends**

**Figure S1:** Raw read counts of 16S sequenced samples across time points and groups.

**Figure S2:** Estimated load of most abundant species, based on read counts as proxy for microbial biomass.

**Figure S3:** 16S read counts plots for negative and positive controls.

**Figure S4:** Tumor area and multiplicity.

**Figure S5:** Tumor growth in transplantable model over time.

#### SUPPORTING DATA

##### Immunohistochemistry (IHC) of immune markers

We performed IHC on CD3, CD4, CD8A, CD11C as well as PD1 and PD-L1 to assess tumor-lymphocyte infiltration and correlate ICI-response with PD1 and PD-L1 expression in a representative set of samples. Our IHC labeling was specific (in experimental samples and positive controls) hence we excluded technical difficulties as an issue. We found that very few T cells were close to or within the pre-malignant and malignant tumor tissue and very few PD1 and PD-L1-positive cells were detected. These results led us to the conclusion that our observations are more likely connected to the hairless (*Hr*) mutant skin phenotype which - particularly in adult mice - has significant abnormalities that result in an influx of T cells and macrophages.

#### SUPPLEMENTARY MATERIALS AND METHODS

##### Animals

The specific-pathogen free (SPF) albino hairless SKH1 mice (Crl:SKH1-$Hr^{hr}$; Strain Code 477, hereafter abbreviated as "SKH1 mice") were obtained from Charles River Laboratories, Wilmington, MA, USA. All mice were housed, and experiments performed in the animal facility at The Jackson Laboratory, Bar Harbor, ME, USA. All experiments were approved by The Jackson Laboratory Animal Care and Use Committees and were performed in accordance with National Institutes of Health regulations. Mice were housed with up to 5 mice per Innovive box.

SKH1 mice (51 mice: 46 females, 5 males) were 6-8 weeks of age at the initiation of UV-B light irradiation. A small number of mice suffered from genital prolapses and were treated with non-antimicrobial lubricants and harvested when treatment failed. The gender imbalance is a result of the sudden deaths of SKH1 mice due to intolerance of antibiotics (see below for more information) in older mice or genital prolapses which resulted in a rearrangement of the study cohort.

##### UV-B treatment

Most mice were UV-B-irradiated Monday, Wednesday, and Friday for a total of 13 weeks. Initially, for 2 weeks, the mice received a total of 750 mJ/cm$^2$ with three 250 mJ/cm$^2$ doses per treatment. For the subsequent 11 weeks, the mice received a total of 1250 mJ/cm$^2$, with two treatments of 500 mJ/cm$^2$ (Mon, Fri) and one treatment of 250 mJ/cm$^2$ (Wed), based on (Chitsazzadeh et al., 2016) and previous inhouse pilot studies. In this mouse 250 mJ/cm$^2$ is equal to 1 minimal erythemal dose (minor sunburn). UV treatment was discontinued after 13 weeks.

Two parallel UVP XX-Series UV Bench Lamp tubes (peak emission at 302nm, UVP34003901, Fisher Scientific) were used with a standard fixture for UV-B-exposure for irradiation from above. Mice were kept in their original cage and the food hopper was replaced with an in-house designed wire mesh lid held by a 3D printed fixture that prevented mice from standing on their hind legs and climbing out of the cage. The UV-B dosage was regularly checked using a calibrated meter (UVB-500C, National Biological Corp., Beachwood, USA).

##### Enrollment

Mice were enrolled into the next phase when they showed at least one lesion ≥4 mm. Mice were mostly randomly assigned to groups. Whenever possible, pairs of mice with similar tumor size were assigned to the immunotherapy subgroup and one to the isotype control subgroup of a treatment group. Mice were enrolled in a staggered manner based on their tumor growth which was assessed by weekly caliper measurements. Enrollments began at 18 weeks after experiment start.

**Antibiotics administration**

All mice received a cocktail of topical antibiotics for 1 week consisting of 1:1 (w/w) Mupirocin (2%, Taro) and Neosporin (bacitracin (400 U), neomycin (3.5 mg), polymyxin B (5,000 U) in petrolatum) twice per week. In parallel, mice received a cocktail of systemic antibiotics consisting of Ampicillin sodium salt (1 mg/ml, Sigma or GoldBio), Vancomycin hydrochloride (0.25 mg/mL, Sigma or GoldBio), Colistin sulfate salt (1 mg/mL, Cayman Chemical or GoldBio), and Streptomycin sulfate salt (5 mg/mL, Sigma or GoldBio) in sterile drinking water. For groups 1 and 2, the mice were supposed to receive systemic antibiotics for 6 days with 1 day of recovery before bacteria/ICI administration. Many mice showed adverse effects to the cocktail (dehydration) and the antibiotics regimen had to be adjusted. We note that this observation was pervasive and specific to the SKH1 mice (also described for other mouse strains by (Almonte et al., 2022)), as C57BL/6J mice did not suffer these effects in similar experiments.

Mice from wave 1 received the original cocktail. After two days they showed signs of dehydration and were allowed one day of recovery with unmedicated water, and then were given a 75% dilution of the cocktail until the end of the week (without exhibiting further signs of dehydration). Mice from wave 2 received the original cocktail for 3 days until showing signs of dehydration, were allowed 1 day of recovery with unmedicated water, and then were given a 75% dilution of the cocktail mixed with a sweetener (5 mg/ml, Truvia Natural Stevia) to make the medicated water more palatable. Since symptoms did not consistently improve, mice were switched to 1mg/ml Ampicillin with sweetener until the end of the week without showing further adverse effects. Mice from waves 4-8 received Ampicillin (1 mg/ml, Sigma) with sweetener only until the end of one week regimen, except for mice assigned to group 3 which received the Ampicillin for 4 weeks in total until harvest (topical antibiotics was applied 2x per week). Mice were under close veterinary supervision and mice with severe dehydration and no signs of recovery were euthanized to comply with the IACUC protocol. Topical antibiotics treatment was not interrupted during the treatment week. Pilot studies with this antibiotics cocktail had not shown adverse effects in younger mice, but these significantly older mice were reluctant to consume antibiotic-treated water, requiring sweetening and adjustments to the cocktail to avoid dehydration. Microbiome depletion was confirmed using 16S sequencing of the skin swab and fecal pellet and cultivation of skin swabs.

**Microbial application**

Upon significant tumor growth (one tumor >4mm$^3$), mice were assigned to one of the groups. Group 1 received topical application of *C. acnes*, Group 2 received oral gavage of *A. muciniphila* while Group 3 received a long-term antibiotics treatment as well as a vehicle control (10% Glycerol in tryptic soy broth (TSB)) topically applied and gavaged. Microbial application/gavage recurred three times per week (Monday, Wednesday, Friday) for 3 weeks after ICI/isotype administration (see below) until harvest.

For both, *C. acnes* (strain 6919, ATCC) and *A. muciniphila* (strain BAA-835, ATCC) were cultured in tryptic soy broth (TSB, Fisher Scientific, DF0370-17-3) for 4 days. CFU (colony forming units) count was assessed using hemocytometers (Fisher Scientific, 22-600-107), cells were condensed by centrifugation, the supernatant decanted, and the pellet resuspended in TSB + 10% glycerol (Sigma, G5516-1L) to obtain a CFU count of 10^8 CFUs per 10µl using sterile technique. Glycerol (Fisher Scientific, BP229-1) was added as a cryoprotectant and as a carbon source for the microbes (Shu et al., 2013). Aliquots were prepared with 220µl per tube for *C. acnes* and 150µl per tube for *A. muciniphila*. Aliquots were frozen at -80°C and dilutions of sample aliquots were recounted to assess accuracy of the CFU counts. Bacteria were also plated on Trypticase™ Soy Agar with 5% Sheep Blood (BBL-TSA II, Fisher Scientific, B21261X), grown under anaerobic conditions and then species identification was confirmed using MALDI-TOF (matrix assisted laser desorption ionization-time of flight, Bruker Daltonics, Germany). Using a sterile transfer device (Puritan, 25-28107), bacteria were directly transferred to a MALDI target spot. We then used the 'extended direct method' for sample preparation, in which 70% formic acid (Sigma-Aldrich, 5438040100) is used to solubilize the bacterial cell wall prior to the addition of matrix (Bruker Matrix HCCA, 14932). One spot on the target was reserved for the bacterial test standard (Bruker, 8255343) for calibration. Mass spectrometry analysis was performed using Flex Control 3.4 software (Bruker Daltonics, Germany).

**Immunotherapy administration**

In parallel to the bacterial treatment, mice received either the anti-PD1 monoclonal antibody (InVivoMAb anti-mouse PD1 (CD279), Clone: 29F.1A12, BioXcell, BE0273, 9.15 mg/ml) or the corresponding isotype control (*InVivo*MAb rat IgG2a isotype control, anti-trinitrophenol, Clone: 2A3, BioXcell, BE0089, 9.22mg/ml). PD1 blockade has been shown to delay SCC development in mice (Belai et al., 2014). Mice were injected intraperitoneally three times per week for 3 weeks with 100µl in PBS (200µg) per dose before bacterial administration.

**Tumor measurements of spontaneous SCC model**

During the experiment, lesions were measured with a caliper, counted, and documented. By gross observation, lesions ≥2mm$^2$ were included in the manuscript as SCC-related lesions. The lesions were included if the length, width, or height of each tumor was 2mm or larger measured at/from the base of the tumor. Measurements were conducted once per week and at the end point. Volume, area and Knud Thomsen's Three-Dimensional (KT-3D) Tumor Surface Area were calculated as described in (Bazin et al., 2020). Tumor multiplicity was determined as the average number of individually countable tumors per mouse. Since mice were enrolled in waves, the data was normalized to the experimental timeline for illustration purposes. Statistical analysis was conducted as described below.

**Skin swabs for cultivation**

A second set of skin swabs for cultivation to assess the CFU counts were taken as above, plated on BBL-TSA II agar plates, and incubated anaerobically for ~4 days at 37°C in 5% $CO_2$. Pictures were taken after incubation for qualitative assessment.

**Skin swabs for DNA extraction**

Skin microbiome dynamics was observed using skin swabs. Dorsal skin swabs were taken on the day of enrollment (TP0) defined by tumor measurements of at least 1 lesion ≥ 4 mm (before antibiotics), after 1 week of antibiotics treatment (TP1), and at the end point (TP2). At the end point, per mouse, 2 skin swab samples were collected: a collective swab of the healthy, tumor-adjacent skin and a collective swab of all lesions. Skin swabs were taken using sterile Puritan™ PurFlock Ultra Flocked Swabs with Elongated Tips (Fisher Scientific, 22-029-506) pre-moistened in sterile phosphate buffered saline or water. Swabs were gently rubbed onto the dorsal skin for 30 seconds, and then stored in 350µl yeast and cell lysis buffer (Epicentre, MPY80200) or Tissue and Cell lysis solution (Epicentre, MTC096H) lysis buffer with 100µl glass beads (0.1 diameter, BioSpec Products, 11079101) at -80°C until shipping. Samples were shipped on dry ice to The Jackson Laboratory for Genomic Medicine, Farmington, Connecticut, USA and stored at -80°C until DNA extraction.

**Fecal sample collection**

Fresh fecal pellets were collected at the same time points as skin swabs (Week 1, Week 2, Week 5). Pellets were placed into RNALater for preservation. Samples were stored at -80°C. Samples were shipped on dry ice to The Jackson Laboratory for Genomic Medicine, Farmington, Connecticut, USA and stored at -80°C until DNA extraction.

**Environmental samples, positive and negative controls**

Environmental samples (fresh mouse chow, bedding) were collected since they consist of plant material and mitochondrial DNA can be amplified and be mistaken as 16S rRNA genes. Samples were extracted as described above. Additionally, per extraction round, one sample of a defined, in-house mock community (25 diverse Gram-positive bacteria, Gram-negative bacteria, and fungi) and a negative reagent control (nuclease-free water, Qiagen) were included. For library generation, a negative control was included as well (nuclease-free water, Qiagen). Library products were measured on the Qubit 2.0 Fluorometer (Thermo Fisher Scientific, Waltham, Massachusetts, USA) and pooled for sequencing (see below).

**Transplantable SKH1 tumor model**

An immortalized cell line was generated from keratinocytes extracted from a histologically confirmed tumor induced using UV-B light in SKH1 mice (similar to above). Cells (clone B610K) were provided by the Tsai Lab (Moffitt Cancer Center, Tampa, Florida, USA). Cells were maintained and passaged in a custom media consisting of 375 ml Gibco™ Ham's F-12 Nutrient Mix (Fisher Scientific 11-765-070), 125mL DMEM with L-Glutamine, Glucose and Sodium Pyruvate (Thermo Fisher, MT10013CV), 20.8 µL Cholera Toxin (0.2mg/mL stock, Sigma, C8052), 250µL Insulin (10mg/mL stock, Sigma, I5500-50MG), 50uL Hydrocortisone (4 mg/mL stock, Sigma, H4881-100MG), 6.5µL Triodo-Thyramine (1.3 µg/mL stock, Sigma, T6397-100MG), 5mL Adenine (2.4 mg/ml stock, Sigma, A3159-5G), 25 ml FBS heat inactivated (5%. VWR SCIENTIFIC INCORPORATED, 500 ml), 5ml L-glutamine (Thermo Fisher, 25030081), 5 ml Pen/strep (Gibco, 15140122), 1ml Primocin™ (VWR, MSPP-ANTPM1).

At every change of media or passage, fresh EGF (10 ng/mL final conc., VWR, 10208-654) was added to the culture and cells were grown in a 5% $CO_2$ incubator at 37°C. Cells were used up to passage 26 for injections. Engraftment efficiency declined with later passages, especially >30 passages.

For injections, the cells were washed in 2x PBS and harvested using Accutase (A6964, Sigma) for about 45 min, spun down at 500g for 10 min and resuspended in 5 ml of warm growth media. The cells were counted and reconstituted in the appropriate amount of 1:1 PBS:Matrigel (Corning, 354230). Insulin syringes (Fisher Scientific, 50-209-2887) with 50µl of cells were prepared and placed on ice until injection. Mice were restrained using a ScruffGuard (Research Devices Ltd., SKU: 000002) and injected intradermally into the right flank. Mice that received Ampicillin treatment via the water as described above (with Truvia) and were treated from the day of the injection until the endpoint (3.5 weeks). Mice who became dehydrated were temporarily removed from the treatment and placed back on normal water for a quick recovery (2-3 days) and then the concentration of Truvia was increased from 5 mg/ml to 7 mg/ml until the end of the experiment.

For the intradermal injection of 250k B610K cells, 10 males with or without (systemic) antibiotics were used. For females, 11 mice on regular water and 9 mice on antibiotics-medicated water were injected. Three females were euthanized due to dehydration, leaving 6 mice at the endpoint. For the injection of 500k cells, 10 males with and 11 without antibiotics were used. Two males on regular water had to be euthanized at week 2.5 due to excessively fast tumor growth, resulting in 9 mice at the week 3.5 endpoint. For females, 11 mice on antibiotics and 8 mice on regular water were used.

**Tumor measurements of transplantable SCC model**

Tumor growth (width and length) was measured weekly and at the endpoint as described above. The ellipsoid volume was calculated with the commonly used formula (Width x Height x Length)/2 (as described in (Tomayko and Reynolds, 1989)), with the assumptions Width = Height and Width < Length. Statistical analysis was conducted as described below.

**Tissue harvest**

At the endpoint, mice were euthanized humanely by cervical dislocation. Mice were sacrificed four days after the last ICI injection. The skin was harvested postmortem and all macroscopically visible lesions including adjacent normal appearing skin were carefully dissected, fixed in neutral buffered 10% formalin overnight, washed in PBS and stored in 70% EtOH until paraffin embedding.

**Histology**

Tissue was paraffin embedded using standard protocols. Blocks were sectioned and H&E stained (using standard protocols) at the Histopathology & Microscopy Sciences at the Jackson Laboratory (Bar Harbor, ME, USA). Lesions were scored blindly by our histopathologist using the criteria described by Benavides et al. and Bazin et al. (Bazin et al., 2020, Benavides et al., 2009). All observations are separately listed in Table S1. Table 2 shows a digest of all histologic observations where fully and microinvasive tumors were combined as "invasive SCC" and preneoplastic and premalignant tumors were combined as "premalignant SCC". Standalone exophytic and fibropapilloma were counted as "papilloma".

**Immunohistochemistry (IHC) of immune markers**

We performed IHC on CD3, CD4, CD8A, CD11C as well as PD1 and PD-L1 on a subset of samples of different SCC severity in order to assess tumor-infiltrating lymphocytes (TILs) and correlate ICI-response with PD1 and PD-L1 expression. All antibody studies were performed on the Leica Bond RX staining instrument using standard protocols at the Histopathology & Microscopy Sciences at the Jackson Laboratory Bar Harbor, ME, USA. For CD3 (ab237707, abcam, 1:500 dilution), CD4 (Thermo-Fisher, 14-9766-80 Rat 1:1000), CD8A (14-0808-80, Thermo-fisher, 1:1000) pretreatment was done with citrate buffer for 20 minutes. For CD11C (ab219799, abcam, 1:100) and PD1 (ab214421, abcam, 1:1000), Tris/EDTA treatment was carried out for 20 minutes. For PD-L1 (ab238697, abcam, 1:500) no pretreatment was performed. As secondary antibody the Rabbit Anti-Rat IgG Antibody (AI-4001, Vectorlabs, 1:500) was used and the labelling was carried out using the BOND Polymer Refine Detection kit (DS9800, Leica).

**DNA extraction and 16S rRNA gene sequencing**

DNA was extracted from skin swabs and fecal pellets using the QIAamp 96 DNA QIAcube HT Kit (Qiagen, 51551) with the following modifications: Before loading on the Qiacube robot, samples were incubated with 5µl Lysozyme (10 mg/ml, Sigma-Aldrich, L6876), 1µl of Lysostaphin (5000U/ml, Sigma-Aldrich, L9043), and 1µl mutanolysin (5000 U/ml, Sigma-Aldrich, M9901) for a digest at 37°C for 30 min. Samples were mechanically disrupted with 100µl glass beads (0.1 diameter, BioSpec Products, 11079101) for 2x 3 min at 30 Hz (TissueLyser II, Qiagen).

16S library preparation was carried out using barcoded primers targeting the V1-V3 region of the 16S rRNA gene (FWD 5'-AGAGTTTGATCCTGGCTCAG-3', REV 5'-ATTACCGCGGCTGCTGG-3'). For amplification (30 cycles at 56°C), the AccuPrime Taq DNA Polymerase High Fidelity and 10X AccuPrime Buffer II (Invitrogen, 12346086) were used. For PCR positive controls, 4 ng of evenly mixed bacteria (HM-782D, HM-276D, BEI Resources) and nuclease-free water was used for PCR-negative controls.

PCR products were purified using the Agencourt AMPure XP Beads (Beckman Coulter, A63882) and quantified using the Qubit dsDNA HS Kit (Thermo Fisher Scientific, Q32854). PCR products were multiplexed to equimolar library pool (final 4 nM), the pool was quantified, and quality checked using the Illumina Library Quantification Kit (ROX Low qPCR Mix, Illumina, KK4873) and Illumina Library Quantification Standards 1-6 (Illumina, KK4903) and on the 4200 TapeStation System (Agilent Technologies, Santa Clara, CA) using the High Sensitivity D1000 ScreenTape Assay (Agilent Technologies, Santa Clara, CA). Sequencing was performed using a 250-bp paired-end sequencing protocol on the Illumina MiSeq platform (Illumina, San Diego, USA) at the Genome Technologies facility at The Jackson Laboratory for Genomic Medicine.

**Quantitative polymerase chain reaction**

Quantitative polymerase chain reaction (qPCR) was carried out on a representative number of skin and fecal samples before and after antibiotics treatment to demonstrate the depletion of the microbiome. The Fast SYBR™ Green Master mix (4385610, Applied Biosystem) and universal 16S primers were used (FWD 5'-TCCTACGGGAGGCAGCAGT-3, REV 5'-GGACTACCAGGGTATCTAATCCTGTT-3, (Nadkarni et al., 2002)) in Fast QPCR plates (C18081-15,

Denville) for qPCR amplification. QPCR was carried out for 40 cycles with 95°C for 1 second and 60°C for 45 seconds on the ViiA™ 7 Real-Time PCR System (Applied Biosystems). The same volume of DNA was used for each sample to compare cycle threshold (Ct) values. Samples were evaluated in triplicate.

**16S rRNA gene data processing**
An in-house civet pipeline was used to quality screen, data cleaning, and classify the sequencing data. Briefly, barcodes and primers were removed from the raw reads using Trimmomatic software (version 0.32 (Bolger et al., 2014)). We removed sequences with low quality (average quality < 35) and ambiguous bases (N's). Paired reads from the same amplicon were assembled and joined using the FLASH assembly software (Magoc and Salzberg, 2011). Chimeric amplicons were removed using the UChime software (Edgar et al., 2011). Reads were further screened against the mouse genome to remove potential host-derived sequences using BMTagger (version: 3.101, (BMTagger, RRID:SCR_014619)). We used USEARCH V11 (Edgar, 2010) to construct Operational Taxonomic Units (OTUs) by grouping of merged cleaned reads that are at least 97% identical. Final taxonomic assignment of OTUs sequences was performed using the Ribosomal Database Project (RDB) classifier (Wang et al., 2007) yielding 781 OTUs.

OTU_1, was the most abundant OTU identified in environmental samples (mouse chow and bedding, Figure S4, insert) and reoccurring in skin and fecal samples. It was subtracted from the dataset as a contaminant from chow and bedding. OTU_1 was classified as *Phyllobacterium* by RDP and confirmed by BLAST. No read number cutoff (usually used as unsupervised feature reduction technique), was applied since we specifically removed and perturbed the microbiome.

All taxonomic features with a mean relative abundance of <0.001% (denoise function, (Arumugam et al., 2011) across the dataset (without controls) and a prevalence of <10% were removed from the dataset. This corrected for potential false positives and for multiple hypothesis testing and yielded 324 OTUs in the skin and fecal samples. Metadata is listed in Table S2 and the full OTU data set is in Table S3.

**Data overview**
A total of 376 samples were sequenced which included 8 in-house mock community samples, of which five were cell mixtures included in the extraction process, and three DNA mixtures of separately extracted bacterial cells. Additionally, one mouse chow and one mouse bedding sample, five extraction negative controls, four PCR negative controls as well as 153 fecal pellet samples and 204 skin swabs were included in the analysis.

**Data analysis and visualization**
R (version 4 (R Core Team, 2020)) and RStudio (version 1.4.94) were used for statistical analysis. Linear regression models (lm R function, R package "stats" (version 3.6.2)) with time and mouse group as interaction terms were fitted, and repeated measure ANOVA (1000 permutations) was used to test statistical significance in tumor burden between two treatment groups over time. P-values were post-hoc multiple-test corrected using the Benjamini-Hochberg method. Corrected p-values <0.05 were deemed significant. To assess whether tumor burden differences were influenced by the gender imbalance we fitted the linear models with all animals and just the females and compared the results. For the UV-induced SCC experiment, weeks 3-5 were included in the analysis (the weeks of ICI treatment). For the transplantable model, all data but week 0 (the week of injection), was included. To assess group differences at individual time points of the transplantable model, the non-parametric (unpaired) Wilcoxon signed-rank test was employed for matched (but independent) comparisons. Differences in tumor severity based on numbers of invasive and premalignant lesions confirmed by histologic evaluation were assessed using Fisher's exact test. R functions used for analysis and data visualization are the following: reshape2, vegan, ggplot2, tidyverse, ggpubr, dplyr, plyr (Kassambara, 2020, Oksanen et al., 2019, Wickham, 2016, 2017, Wickham et al., 2019).

#### SUPPLEMENTAL TABLE LEGENDS

**Table S1: Detailed list of histologic observations.** Based on sum observations across mice of each group. Numbers in brackets are observations for female mice only for the groups that included 1-2 male mice.

| | 1A | 1B | 2A | 2B | 3A | 3B |
|---|---|---|---|---|---|---|
| | 8 mice | 9 mice | 9 mice | 9 mice | 8 mice | 8 mice |
| **Spindle anaplastic sarcoma** | 0 | 4 | 3 (0) | 2 (2) | 0 | 0 |
| **Keratoacanthoma** | 0 | 0 | 0 | 0 | 0 | 2 (2) |
| **Fully invasive SCC** | 3 | 4 | 2 (1) | 3 (2) | 1 (1) | 0 |
| **Microinvasive grade 3** | 2 | 0 | 0 | 0 | 1 (1) | 1 (1) |
| **Malignant microinvasive grade 3/ Fibropapilloma** | 0 | 1 | 0 | 0 | 0 | 0 |
| **Microinvasive grade 3/trichoepithelioma** | 0 | 0 | 0 | 2 (2) | 0 | 0 |
| **Microinvasive grade 2** | 2 | 1 | 3 (2) | 2 (2) | 0 | 0 |
| **Microinvasive grade 1** | 3 | 5 | 6 (6) | 4 (1) | 3 (3) | 0 |
| **Microinvasion grade 1 with Fibropapilloma** | 0 | 0 | 1 (1) | 0 | 0 | 0 |
| **Malignant microinvasive grade 1/ trichoepithelioma** | 0 | 0 | 1 (1) | 0 | 0 | 0 |
| **Preneoplastic grade 2/ exophytic papilloma** | 0 | 0 | 0 | 2 (2) | 1 (1) | 2 (2) |
| **Preneoplastic grade 2/ trichoepithelioma** | 0 | 0 | 0 | 0 | 1 (1) | 0 |
| **Preneoplastic grade 1** | 0 | 0 | 0 | 0 | 1 (1) | 0 |
| **Premalignant grade 3** | 1 | 2 | 3 (3) | 2 (1) | 1 (1) | 1 (1) |
| **Premalignant grade 3/ exophytic papilloma** | 2 | 0 | 1 (1) | 0 | 0 | 0 |
| **Premalignant grade 3/ mixed trichoepithelioma & papilloma** | 0 | 1 | 0 | 0 | 0 | 0 |
| **Premalignant grade 3/ trichoepithelioma** | 1 | 2 | 0 | 0 | 0 | 0 |
| **Premalignant grade 2** | 7 | 4 | 4 (4) | 2 (2) | 4 (4) | 14 (14) |
| **Premalignant grade 2/mixed** | 1 | 0 | 1 (1) | 0 | 0 | 0 |
| **Premalignant grade 2/exophytic papilloma/ trichopapilloma** | 0 | 0 | 0 | 0 | 0 | 0 |
| **Premalignant grade 2/exophytic papilloma** | 14 | 6 | 9 (3) | 7 (7) | 4 (2) | 6 (2) |
| **Premalignant grade 2/trichoepithelioma** | 7 | 5 | 4 (2) | 4 (4) | 0 | 2 (2) |
| **Premalignant grade 1/2, exophytic papilloma** | 0 | 0 | 2 (0) | 0 | 2 | 0 |
| **Premalignant grade 1** | 4 | 8 | 3 (3) | 2 (1) | 8 (6) | 0 |
| **Premalignant grade 1/exophytic papilloma** | 2 | 1 | 2 (2) | 0 | 0 | 0 |
| **Premalignant grade 1/trichoepithelioma** | 1 | 1 | 3 (2) | 1 (1) | 1 (1) | 2 (1) |
| **Premalignant grade <1** | 0 | 0 | 0 | 2 (2) | 0 | 1 (1) |
| **Fibropapilloma** | 0 | 0 | 0 | 0 | 0 | 0 |
| **Exophytic papilloma** | 0 | 1 | 1 (1) | 0 | 0 | 0 |
| **Reactive lymphnode** | 0 | 2 | 0 | 0 | 0 | 0 |
| **Telangectasia with thrombus** | 0 | 0 | 1 (1) | 0 | 0 | 0 |
| **Acanthosis/ FB granuloma / normal Hr phenotype** | 3 | 7 | 5 (5) | 4 (2) | 4 (4) | 5 (5) |
| **Ulcer** | 0 | 5 | 4 (3) | 2 (2) | 1 (1) | 1 (1) |
| **Cyst** | 1 | 0 | 1 (1) | 1 (1) | 4 (3) | 0 |
| **Erosion** | 0 | 2 | 1 (1) | 2 (2) | 0 | 0 |
| **Invasive trichoepithelioma grade 3** | 0 | 0 | 0 | 0 | 0 | 1 (1) |
| **Trichoepithelioma** | 0 | 1 | 1 (1) | 0 | 0 | 0 |
| **Pseudoepitheliomatous hyperplasia** | 0 | 0 | 0 | 0 | 1 (1) | 0 |
| **NA** | 1 | 1 | 0 | 0 | 0 | 4 (4) |

**Table S2: Metadata.** Metadata for each sample collected and included in the study.

- See separate excel table

**Table S3: 16S read counts table.** The table includes all 376 samples and 781 OTUs (before curation).

- See separate excel table

**Table S4: Table of all p-values calculated for each comparison.** Linear regression models with time and mouse group as interaction terms were fitted and repeated measure ANOVA. Highlighted in green are statistically significant after multiple test correction ($p<0.05$). Details about the calculation of the parameters are described in the Supplementary Material and Methods. ICI: Immunotherapy, Abx: Antibiotics.

| | **Group Comparison** | **KT-surface** | | **Multiplicity** | | **Total area** | | **Volume** | |
|---|---|---|---|---|---|---|---|---|---|
| | | **All mice** | **Females only** | **All mice** | **Females only** | **All mice** | **Females only** | **All mice** | **Females only** |
| **ICI vs. Isotype control** | **1A vs 1B** | 0.099 | 0.077 | 0.206 | 0.195 | 0.046 | 0.048 | 0.376 | 0.346 |
| | **2A vs 2B** | 0.244 | 0.236 | 0.877 | 0.873 | 0.021 | 0.044 | 0.029 | 0.207 |
| | **3A vs 3B** | 0.003 | <0.001 | 0.735 | 0.726 | 0.001 | <0.001 | <0.001 | 0.001 |
| **Bacteria vs. Abx** | **1A vs 3A** | 0.013 | 0.019 | 0.111 | 0.158 | 0.025 | 0.036 | 0.013 | 0.025 |
| | **1B vs 3B** | 0.083 | 0.11 | 0.045 | 0.047 | 0.113 | 0.176 | 0.161 | 0.267 |
| | **2A vs 3A** | <0.001 | 0.006 | 0.183 | 0.876 | 0.003 | 0.037 | 0.001 | 0.012 |
| | **2B vs 3B** | 0.162 | 0.654 | 0.597 | 0.753 | 0.25 | 0.982 | 0.1 | 0.497 |

#### SUPPLEMENTAL FIGURE LEGENDS

**Figure S1: Raw read counts of 16S sequenced samples across time points and groups**. Boxplots show total microbial read counts identified per sample in **A**: skin and **B**: fecal samples before (week 1, wk1) and one week after (week 2, wk2) antibiotics treatment as well as at the end point (week 5, wk5) for each group. HS: healthy skin and TU: tumor at endpoint. Metadata is listed in Table S2, and the Operational taxonomic unit (OTU) table is in Table S3. Depicted is the data for all mice irrespective of sex: Group 1A: n=8, Group 1B: n=9, Group 2A: n=9, Group 2B: n=9, Group 3A: n=8, Group 3B: n=8.

**Figure S2: Estimated load of most abundant species, based on read counts as proxy for microbial biomass**. Abundance plot **A** shows the microbial communities in skin and **B** in the fecal samples before (week1, wk1) and after one week (week2, wk2) of antibiotics treatment as well as at the end point (week 5, wk5). Microbes that were applied topically or gavaged in the respective groups were highlighted in red (*C. acnes*) and cyan (*A. muciniphila*). HS: healthy skin and TU: tumor at endpoint. Metadata is listed in Table S2 and the OTU table is in Table S3. Depicted is the data for all mice irrespective of sex: Group 1A: n=8, Group 1B: n=9, Group 2A: n=9, Group 2B: n=9, Group 3A: n=8, Group 3B: n=8.

**Figure S3: 16S read counts plots for negative and positive controls**. Depicted are the 25 most abundant OTUs identified in the control samples while the remainder OTUs are summarized as "Bacteria". An in-house mock community (25 diverse Gram-positive bacteria, Gram-negative bacteria and fungi) was used as positive controls. The insert shows the enlarged counts plot for bedding and food. Metadata is listed in Table S2 and the OTU table is Table S3. OTU1, a *Phyllobacterium* was identified in bedding and food controls and removed from the dataset (see Material and Methods).

**Figure S4: Tumor area and multiplicity**. **A** and **C**: Depicted are the tumor area and (**B** and **D**) tumor multiplicity for the *C. acnes* (Group 1, **A**, **B**) and *A. muciniphila* (Group 2, **C**, **D**) treated groups compared to the antibiotics treated mice (Group 3). Week 0 represents the initial week of tumor measurement (enrollment), week 1 pre-antibiotics treatment and weeks 2-5 bacteria/antibiotics as well as ICI/isotype treatment with euthanasia in the final week (end point, week 5). P-values were calculated using linear modeling and repeated measure ANOVA including the weeks of treatment 3, 4 and 5 (details in Material and Methods and full list of p-values in Table S4). Data of all mice per group are shown irrespective of sex: Group 1A: n=8, Group 1B: n=9, Group 2A: n=9, Group 2B: n=9, Group 3A: n=8, Group 3B: n=8 (Table 1).

**Figure S5: Tumor growth in transplantable model over time**. Boxplots show the volume of the tumors transplanted into female (**A** and **C**) and male (**B** and **D**) mice injected with different cell numbers (250k and 500k per injection) with and without antibiotics. Tumor growth was significantly reduced in male mice treated with antibiotics versus untreated mice when 250k cells were transplanted. Group differences were assessed using repeated measure ANOVA. Initially, 8-11 mice per group and sex were used but a few mice dropped out during the course of the experiment due to adverse side effects (such as dehydration, detailed in Material and Methods). ns: not significant.
